# A quantitative portrait of habituation in *Stentor coeruleus*

**DOI:** 10.64898/2026.06.09.731162

**Authors:** Tejas Ramdas, Nhi Doan, Austen Theroux, Samuel J. Gershman

## Abstract

Habituation—the decrement in response to a series of stimuli—is a widespread form of learning observed across many organisms, including the unicellular organism *Stentor coeruleus*. A lesser-known feature of *Stentor* habituation, shared with animals, is potentiation: faster habituation to a second stimulus series despite partial or complete recovery of responsiveness before that series begins. This form of metaplasticity suggests that although the first-order habituation memory can decay during the recovery period between the two series, a persistent second-order memory mediates faster relearning. We investigate the response profile of *Stentor* across a range of stimulation frequencies and recovery periods to identify the timescales at which these memory traces operate. We introduce a statistical framework to infer both population and single-cell learning parameters, allowing us to quantify prior qualitative findings and examine relationships among parameters across cells. Two key findings are that potentiation is frequency-sensitive, and that recovery and potentiation are decoupled, consistent with a serial and hierarchical cascade of leaky integrator units underlying these processes. This quantitative portrait provides a foundation for mechanistic modeling of intracellular memory in *Stentor*.

## Introduction

Learning is often viewed as something done by brains, or brain-like systems such as artificial neural networks. However, evidence suggests that learning is ubiquitous across the tree of life, including protozoa (***Gershman et al., 2021***; ***Tang and Marshall, 2018***; ***Dussutour, 2021***), fungi (***Ortega and Gamow, 1970***), and plants (***Abramson and Chicas-Mosier, 2016***; ***Gagliano et al., 2016***). This evidence raises two important questions. First, what forms of learning are these systems capable of? Second, what mechanisms implement learning in these systems?

One possibility is that non-neural systems are only capable of “simple” forms of learning, with more “sophisticated” forms requiring brains. However, even apparently simple forms of learning are more complex than they seem at first glance. Habituation, the response decrement resulting from repeated stimulation, is a good case in point. It is observed in virtually every organism where it has been studied. In many animal species, it is known to exhibit a set of hallmark properties (***Rankin et al., 2009***). The complete set of properties defies explanation in terms of simple mechanisms.

Non-neural systems could be considered sophisticated learners if they exhibit the same hallmark properties. ***Wood*** (***1969***) found evidence for many of these properties in the unicellular organism *Stentor coeruleus*. These cells live in ponds, where they anchor to the floor and filter-feed through an oral apparatus. When threatened (e.g., by perturbing them with a mechanical stimulus), they contract into a sphere, or detach and swim away if the threat is sufficiently severe. This defensive response prevents feeding; the cells thus have an incentive to tolerate mild disturbances.

The severity of threat may be ambiguous: does the disturbance reflect harmless water turbulence or the attack of a predator? Since *Stentor* cannot know the answer *a priori*, it needs to learn from experience. Habituation reflects this learning process: the cell stops contracting in response to a repeated mild disturbance. ***Wood*** (***1969***) showed that *Stentor* habituation goes beyond simple response decrement. It also exhibits *potentiation*, another property observed in animals (***Konorski, 1948***; ***Thompson and Spencer, 1966***): if cells are exposed, after a recovery period, to a second stimulation series, their contraction probability returns to baseline, but then decreases more rapidly than in the first series. In other words, relearning is faster than learning. This implies that *Stentor* has a form of long-term “inactive” memory (***Lewis, 1979***) that can persist silently even after short-term habituation has waned. ***Rajan and Marshall (2025***) recently replicated the potentiation effect in *Stentor*.

Despite this body of evidence, potentiation remains poorly understood. Our contribution is to address several open problems. First, the definition of potentiation is not always clear in past studies. There are several ways to characterize learning speed which have different theoretical implications. One of our contributions is a multi-faceted quantitative definition that allows us to evaluate several theoretical proposals based on empirical data.

A second problem is that a quantitative approach requires data collection at a larger scale than in past studies, due to the stochastic nature of *Stentor* contraction (***Rajan et al., 2023***). Past studies have relied on sample sizes on the order of several dozen cells, but we find that reproducible quantitative results require sample sizes on the order of several hundred. We report large-scale data for several parametric manipulations, allowing us to address fine-grained questions about the structure of learning in *Stentor*.

A third problem, recognized by ***Rajan et al. (2023***), is that analysis of population-level results can be misleading about learning processes at the level of single cells. For example, gradual learning curves at the population level may be artifacts of averaging many abrupt learning curves at the single-cell level (cf. ***Gallistel et al., 2004***). We develop a probabilistic modeling framework for characterizing learning at both the single-cell and population levels. This allows us to measure the statistical evidence for various quantitative hypotheses, further constraining theoretical models. In particular, our results indicate that learning mechanisms operating at multiple timescales are necessary to explain the structure of habituation and potentiation in *Stentor*.

### Adjudicating between computational models of habituation

A computational study of habituation requires a precise definition of the response profile. In the context of potentiation and other “savings” phenomena, experimental studies often define learning speed in different ways, making these phenomena challenging to formalize in a consistent way. One canonical view of learning speed in the psychology literature is the number of trials/stimuli to acquisition of some arbitrary behavioral threshold (***Gallistel and Gibbon, 2000***). In the context of habituation, this behavioral threshold could be an absolute response probability, a relative response probability (fraction of initial response), or a proportional change (fraction of previous response). In what follows, we will be explicit about which definition of learning speed we (or other researchers) are using.

A classic mechanistic model of habituation is based on the leaky integrator, a simple first-order linear ordinary differential equation (***Staddon, 1993***). A special case of this motif (a “perfect” integrator) has also been mapped onto the signaling network underlying adaptation in bacterial chemotaxis, highlighting its utility as an intracellular information-processing architecture (***Alon et al., 1999; Yi et al., 2000***). Computational models of habituation make distinct mechanistic claims about the organization of leaky-integrator motifs underlying the characteristics of habituation (***Staddon, 1993***).

***Staddon*** (***1993***) demonstrated that a single leaky integrator is sufficient to account for one aspect of frequency sensitivity: faster learning (defined in terms of relative response probability) under higher-frequency stimulation. However, reproducing the complementary observation that recovery proceeds more rapidly following higher-frequency stimulation requires a cascade of leaky integrators with appropriately tuned time constants. Distinguishing between these categorically different circuit topologies therefore requires quantitative measurements of both learning curves and recovery dynamics across a range of stimulus frequencies and recovery intervals.

Recent modeling work has sought to extend these motifs to study potentiation. ***Smart et al. (2024***) argue that a single leaky integrator displays potentiation, although they do not have an explicit definition of faster learning. ***Eckert et al. (2024***) present a range of leaky integrator models with different topologies and argue that they display potentiation by demonstrating that they take fewer stimuli to achieve some proportional change. The main issue with prior analyses of potentiation is that it’s unclear if faster learning is simply a result of imperfect recovery, since these deterministic dynamical systems can only return to baseline asymptotically, or if it’s the result of an independent process. Inspired by ***Yi et al. (2000***), we examine potentiation in the context of this dynamical system more rigorously and propose an alternative analysis that obviates this issue.

### Statistical analysis of single-cell habituation profiles

Building on this foundation of experimental and computational studies of habituation, including in *Stentor*, our study seeks to fill in gaps and make claims about the computational motifs and their topologies underlying habituation at the intracellular level. Our experimental paradigm consists of two sequences of stimuli, each of which we refer to as a trial, separated by a recovery period (the intertrial interval, or ITI). We combinatorially manipulate two key stimulus parameters: the inter-stimulus interval (ISI), which determines the frequency of stimulation, and the ITI, to construct a comprehensive quantitative portrait of the habituation profile of *Stentor*.

A population-level analysis, as typically employed in habituation studies, does not suffice to understand the mechanistic basis of habituation at the level of individual cells. Moreover, the behavior of an individual *Stentor* cell is binary (contraction or no contraction) and stochastic, which makes studying single-cell behavior challenging. We collected data at the single-cell level across all conditions and developed a statistical framework to extract the latent probability curves that correspond to the output of the putative intracellular mechanism. Our hierarchical probabilistic framework not only allows us to extract gross population-level estimates of key features of the single-cell habituation curves, but also allows us to compare the joint distributions of parameters across individuals. This allows us to quantify coupling between these parameters. Lastly, we can convert the single-cell habituation curves into phase portraits that show the relationship between response level and learning rate; these phase portraits are informative about computational mechanisms, as discussed below.

With these data and tools in hand, we below identify relationships between responsiveness, learning rate, recovery, and potentiation in *Stentor* habituation; re-examine claims about the frequency-sensitivity of the learning rate; report new findings about the frequency-sensitivity of recovery; and quantify the timescales over which these processes operate. This provides clues about both the classes of models and the rate constants underlying *Stentor* habituation, as has been done in PC12 cells (***Cheever and Koshland, 1992***).

## Results

### Defining potentiation

All of the studies we consider have the following structure. A stimulus is repeated ***N*** times, separated by a regular ISI. The stimulus frequency is defined as 1/ISI. A response is measured for each stimulus number (the repetition index); we will sometimes refer to this index simply as the “stimulus.” Because responses may be stochastic, the object of interest is the response distribution (or some summary statistic such as the mean) over the course of the sequence—the *learning curve*. For binary responses (such as the contraction response studied here), the mean response is equivalent to the response probability. We will refer to one stimulus sequence as a *trial*. After the first trial is completed, an ITI is interposed before the second trial. We will sometimes refer to the ITI as the *recovery period*, because the mean response at the beginning of the second trial approaches the mean response at the beginning of the first trial. We will refer to the mean response at the beginning of each trial as the *baseline*, and the mean response at the end of each trial as the *asymptote* (note that we have chosen *N* to be large enough to ensure that the response probability plateaus by the end of each trial).

If one compares the mean response to stimulus number *s* across the first and second trials, a standard finding is that the response is lower on the second trial, except for the first stimulus. This pattern is often taken to indicate potentiation; we will refer to it as “weak” potentiation to distinguish it from “strong” potentiation, defined below. Note that this definition requires a restriction of the ITI to values that are neither very short nor very long. If the ITI is very short, the mean response may not have recovered back to baseline when the second trial starts. If the ITI is very long, the learning curve may look the same for the first and second trials, indicating that the habituation memory has been lost.

The problem with comparing mean responses directly across trials is that this confounds several conceptually distinct properties of the learning curves. A difference between trials could appear for any of the following reasons: (i) a difference in baseline response; (ii) a difference in the speed of learning; (iii) a difference in the asymptote of learning. We contend that potentiation is properly defined as a difference in the speed of learning. To disentangle this property from baseline and asymptote effects, the relevant measure is the *learning rate*—i.e., the rate of change in the mean response. In a later section, we will describe a probabilistic model with formal definitions of baseline, asymptote, and learning rate.

### Beyond potentiation: phase portraits and second-order memory

Potentiation is one example of a more general phenomenon, what we will call *second-order memory*. To explain this, we will first define *first-order memory*. Let *y* denote the mean response and *u* denote the stimulus input. A general way to describe simple habituation (the response decrement with repeated stimulation) is via an ordinary differential equation (***Smart et al., 2024***):

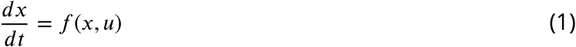

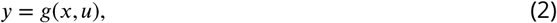

where *f* (*x, u*) is a dynamics function describing how a state (or memory) variable *x* changes with time and stimulus input. The transfer function *g*(*x, u*) describes how the state variable is mapped to the mean response. We will refer to *x* as a first-order memory because its first-order dynamics encode information about the stimulus history. Following ***Smart et al***. (***2024***), we model recovery by assuming that *x*(*t*) → 0 asymptotically in the absence of a stimulus (*u* = 0). In this limit, all information about stimulus history is erased.

For a given trial, the learning rate may change across stimulus number; as we show below, learning is often slowest at the beginning and end of a trial. Moreover, the learning rate may vary across experimental conditions (e.g., different choices of ISI and ITI) even for the same stimulus number in the same trial. We suggest that much of this variability can be captured by a *phase portrait*—the learning rate as a function of the mean response. Because the mean response almost always declines monotonically with stimulus number, the phase portrait captures the relationship between learning rate and stimulus number. In addition, it captures important aspects of the relationship between learning rate and temporal parameters (ISI, ITI) through their effect on the mean response. Intuitively, any effects on the baseline and asymptote should be reflected in the mean response, so that analyzing the learning rate as a function of mean response factors out these confounds.

The phase portrait allows us to define a strong form of potentiation as any difference between the phase portraits for the first and second trials. It is strong in the sense that it places a more stringent requirement on the structure of memory than the weak form, as we discuss next.

### Strong potentiation requires a second-order memory

A classic model of habituation is the discrete-time integrator with a non-linear output function (***Staddon, 1993***; ***Staddon and Higa, 1996***). By chaining these units together, Staddon and colleagues showed that a wide range of habituation phenomena could be captured, including the frequency-sensitivity of habituation and recovery. This idea has served as the basis of biochemically plausible implementations (***Eckert et al., 2024***).

***Smart et al. (2024***) have sought to characterize the simplest mechanistic model consistent with the hallmarks of habituation, including potentiation. Their minimal model takes the form given above in equations 1 and 2. We will refer to the state variable *x* as a “first-order” memory to highlight its direct dependence on the input, and contrast it with “second-order” memories that are mediated by first-order memories.

The transfer function *g*(*x, u*) governs how the memory variable interacts with the input to produce the output. For example, ***Smart et al. (2024***) suggest the Hill function *g*(*x, u*) = ^1^with parameter (Hill coefficient) *n ≥* 1, which constrains the response between 0 and *u*. As long as *x* does not return completely to the baseline, this system will show faster learning in the second trial due to the non-linearity (i.e., weak potentiation). However, it *cannot* show faster learning if it completely returns to baseline (i.e., no strong potentiation). To prove this fact, we construct the phase portrait by expressing the output dynamics as a function of *y* (leaving the dependence on *u* implicit):

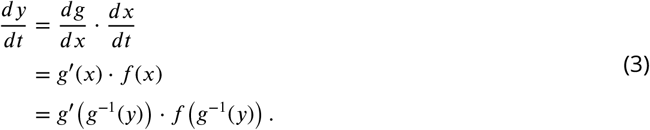

This shows that the phase portrait cannot shift across trials without changing *u, f* , or *g*. However, *u* is identical across trials by design, while *f* and *g* are presumably fixed structurally (absent any evidence to the contrary). Note that we have assumed here invertibility of the transfer function. This assumption does not hold if the transfer function is constant over some range of *x* (e.g., if *g* is a step function, which would obtain in the limit *n* → ∞).

One way to allow the phase portrait to shift acr→oss trials is to introduce a second-order memory (*z*):

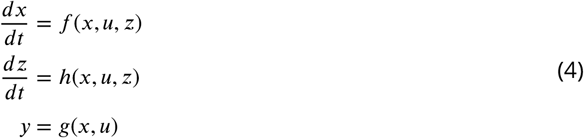

Thus, a second-order memory is sufficient for producing strong potentiation, as long as it modulates the dynamics and transfer functions in such a way that learning is faster on the second trial. Our experiments, described next, are designed to measure evidence for first-order and second-order memories in *Stentor* mechanosensory habituation.

### Experimental setup

We designed an apparatus consisting of a solenoid that taps a dish containing *Stentor* from below, as shown in Figure 1. The anchored cells contract in response to the mechanical force. Consistent with Wood’s experiments, a single “trial” consists of a sequence of 60 stimuli separated by a fixed ISI. The entire experimental protocol consists of an initial trial, followed by an ITI, followed by a second trial with the same ISI as the first. Our automated setup allows us to vary the ISI and ITI as desired. We imaged the cells from above and annotated the contractions of the individual cells that remained anchored through both trials, also consistent with Wood’s protocol. A condition refers to a combination of ISI (1, 2, or 3 minutes) and ITI (1, 2, 3, or 5 hours). This yields a sequence of 120 binary responses for each of 100 cells per condition, resulting in 144,000 binary responses across all 12 conditions. Our experimental paradigm reproduces the basic habituation and potentiation results from prior work. *Stentor* habituate to the sequence of 60 stimuli in the first trial, recover toward baseline in the intertrial interval, and habituate faster in the second trial (Figure 1D).

**Figure 1.**
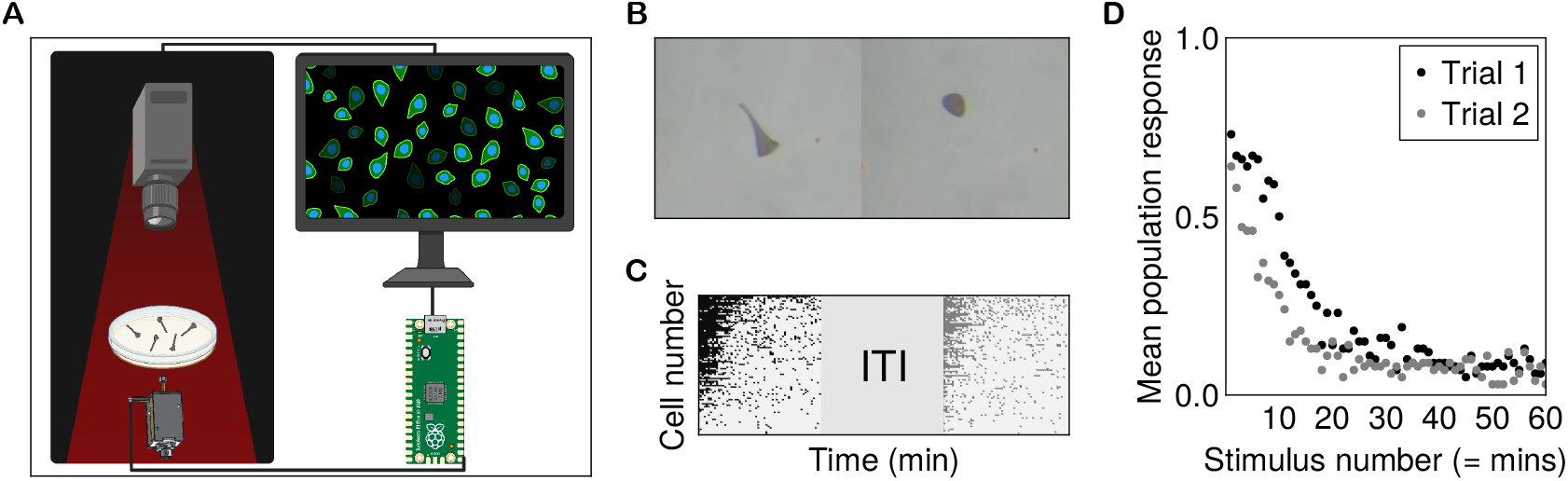
Experimental setup for studying single-cell-level habituation and potentiation in *Stentor*. **(A)**Experimental apparatus: 20–30 *Stentor* cells are placed in a dish above a solenoid that delivers taps. Overhead video recordings are analyzed to extract contraction events. **(B)** Image showing a single *Stentor* extended before a tap and contracted after the tap. **(C)** The experimental protocol consisted of a first trial with a sequence of mechanical stimuli at a fixed interstimulus interval (ISI), followed by an intertrial-interval (ITI), followed by a second trial identical to the first. This yields a set of binary contraction responses for each cell at each stimulus during each trial. **(D)** Population mean response as a proxy for the latent contraction probability. Cells habituate to the series of stimuli, recover close to their baseline response level after a recovery period, and habituate faster in the second trial compared to the first.

### Statistical inference of single-cell habituation profile

We built a hierarchical probabilistic model for single-cell behavior using the Turing software package (***Fjelde et al., 2025***), with parameters inferred from the single-cell contraction responses. The architecture of this model is described in Box 1. We chose to model the habituation of the contraction probability of a single *Stentor* cell as a function of stimulus number with a Hill function. The motivation is twofold: the Hill function allows us to interpolate between gradual and sigmoidal habituation profiles, and Hill functions are commonly used to model the dose-response curves of enzymes and receptors.

We parameterized the habituation function for each trial with four parameters that govern the shape of the function and allow us to extract key properties of the response profile: the Hill coefficient, which governs switchiness (*n*); N50, which reflects the number of stimuli to half-max (*k*); the initial response probability (*i*); and the asymptotic response probability (*a*). The hierarchical form of this model gives us both population-level estimates for these single-cell curve parameters and estimates for each individual cell. The response profile is constrained to be smooth and monotonic, as we would expect of the underlying memory variable. The response probability *y* for cell *j* at stimulus ***s*** is thus:

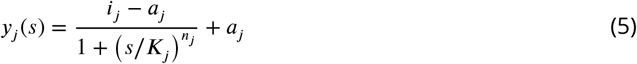

The putative molecular species (e.g., intracellular calcium concentration) controlling contraction probability is continuous in time, but we can only probe it at discrete points. Consequently, *y* is continuous in time, but we reparameterize ***s*** to refer to the time point *t* at which the corresponding stimulus occurs. *y* interpolates the contraction probability between the stimulus time points using the Hill function.

This parameterization allows each of the key features of the habituation curve to vary independently. The Hill coefficient parameter (*n*) allows us to determine the extent to which a single cell follows a switch-like response without performing a separate test for step-like behavior. A high value (*>* 1) suggests a switchy habituation profile, whereas a low value (*<* 1) suggests a gradual habituation profile. The N50 parameter is particularly relevant to potentiation because it governs how quickly the curve reaches its half-max value. A lower N50 value implies faster learning. Thus, the change in this parameter between trials 1 and 2 should reflect potentiation. The initial response and final response parameters allow us to measure the overall responsiveness of the cell and factor it out from the N50 parameter. As such, we should in principle be able to extract the learning rate independent of the response probability, since the N50 parameter is defined in relative rather than absolute terms.

We fit our model independently to each experimental condition (see Materials and Methods for more details). Each cell has 60 responses per trial, yielding 120 binary values. Our model summarizes these values into interpretable parameters for each cell in each trial from which the inferred latent probability curve can be reconstructed using the Hill function. Figure 2 illustrates that our model can successfully summarize the raw single-cell contraction data for the 1 min ISI, 1 hr ITI condition. To verify this beyond visually inspecting Figure 2B, we recomputed the inferred population mean by averaging the response probabilities across all cells at each stimulus, analogous to calculating the population mean from the binary contraction data. Our model recovers the population mean while also exposing cell-to-cell variability. There is considerably more variability in the second trial due to heterogeneity in recovery times. The single-cell curves decrease much more sharply than the population mean would suggest, while the population mean decreases more gradually due to heterogeneity in half-max times. This is consistent with the finding that single cells appear to habituate in a switch-like manner to a 1-minute ISI trial (***Rajan et al., 2023***).

**Figure 2.**
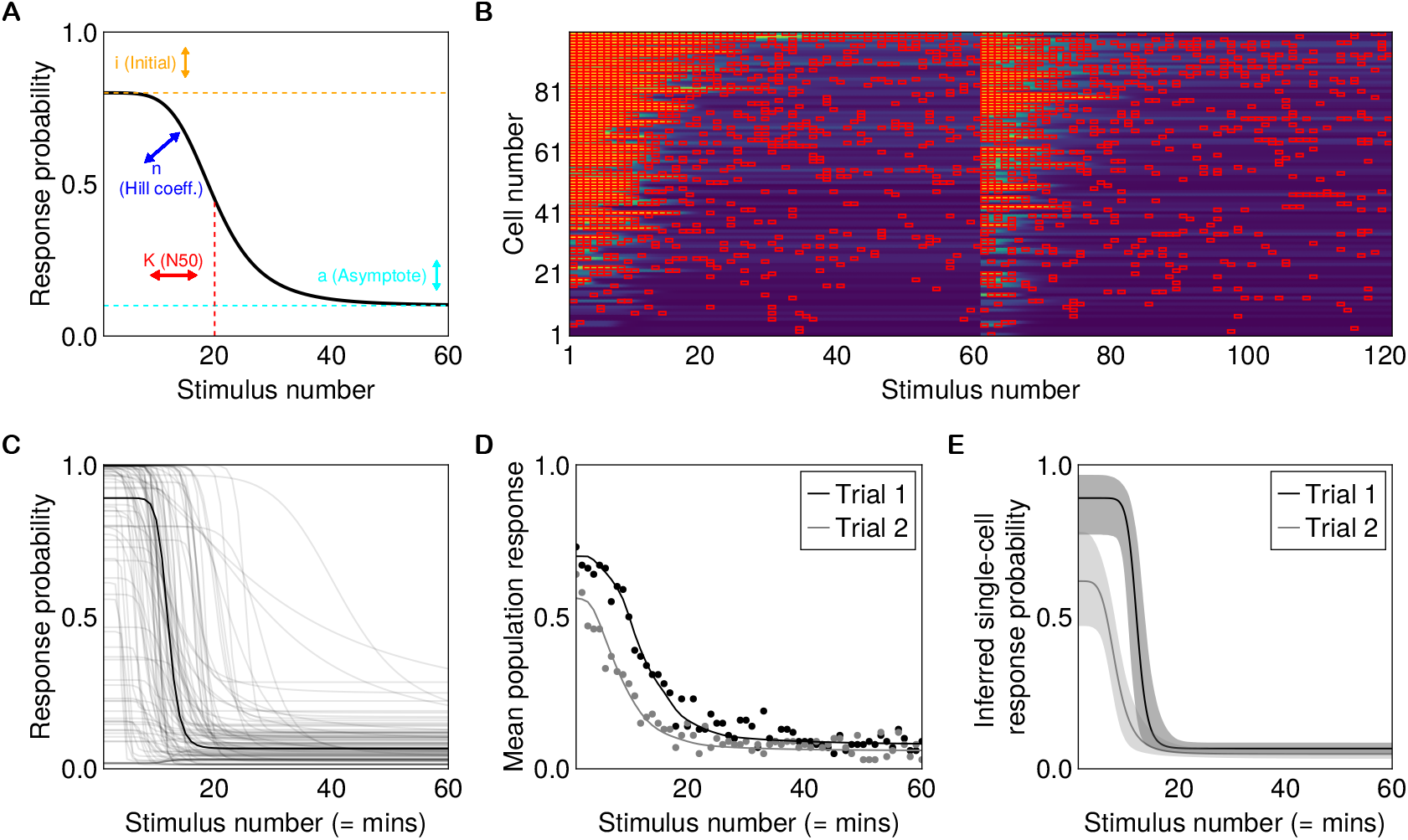
Statistical model recovers naive analysis of response probability. **(A)** Schematic showing the parametrization of the Hill function. Each parameter controls a corresponding aspect of curve shape. **(B)** Each row shows the inferred latent response probability for each cell at each stimulus. Cells have been sorted by cumulative response probability across both trials. **(C)** We obtain a unique latent probability function for each cell that can be averaged to get a point estimate at the population level. **(D)** The mean of the single-cell response curves (solid) faithfully recovers the raw population mean (dotted). **(E)** Single-cell curves, plotted with 95% credible interval bands, decrease much more sharply than the population mean would suggest. The population mean decreases more gradually due to heterogeneity in half-max times.

### Several loosely coupled processes underlie *Stentor* habituation

Having established the validity of our model, we quantitatively examine the relationship between response-curve parameters at the population level and across cells in Figure 3 for the 1 min ISI, 1 hr ITI condition. First, we examine the claim that *Stentor* habituate in a switch-like manner, particularly to 1 min ISI stimuli, as suggested by Rajan et al. We find that the posterior density of the Hill coefficient is heavily concentrated around values significantly greater than 1 (median 11.6, 95% credible interval 6.7–20.5). The single-cell curve is thus much sharper than the population mean, supporting the idea of a switch-like transition. If habituation is due to modification of mechanoreceptors by an intracellular process, this interaction is likely to be highly cooperative. Our results support the finding that *Stentor* decrease their response probability almost instantaneously (***Rajan et al., 2023***). However, the minimum time resolution is approximately 1 minute, given the lower bound on the ISI imposed by re-extension time. As ***Wood*** (***1969***) suggested, it is difficult to study habituation at higher frequencies due to issues with double-counting. Thus, even this putatively instantaneous process could take on the order of minutes to modify the mechanoreceptor.

**Figure 3.**
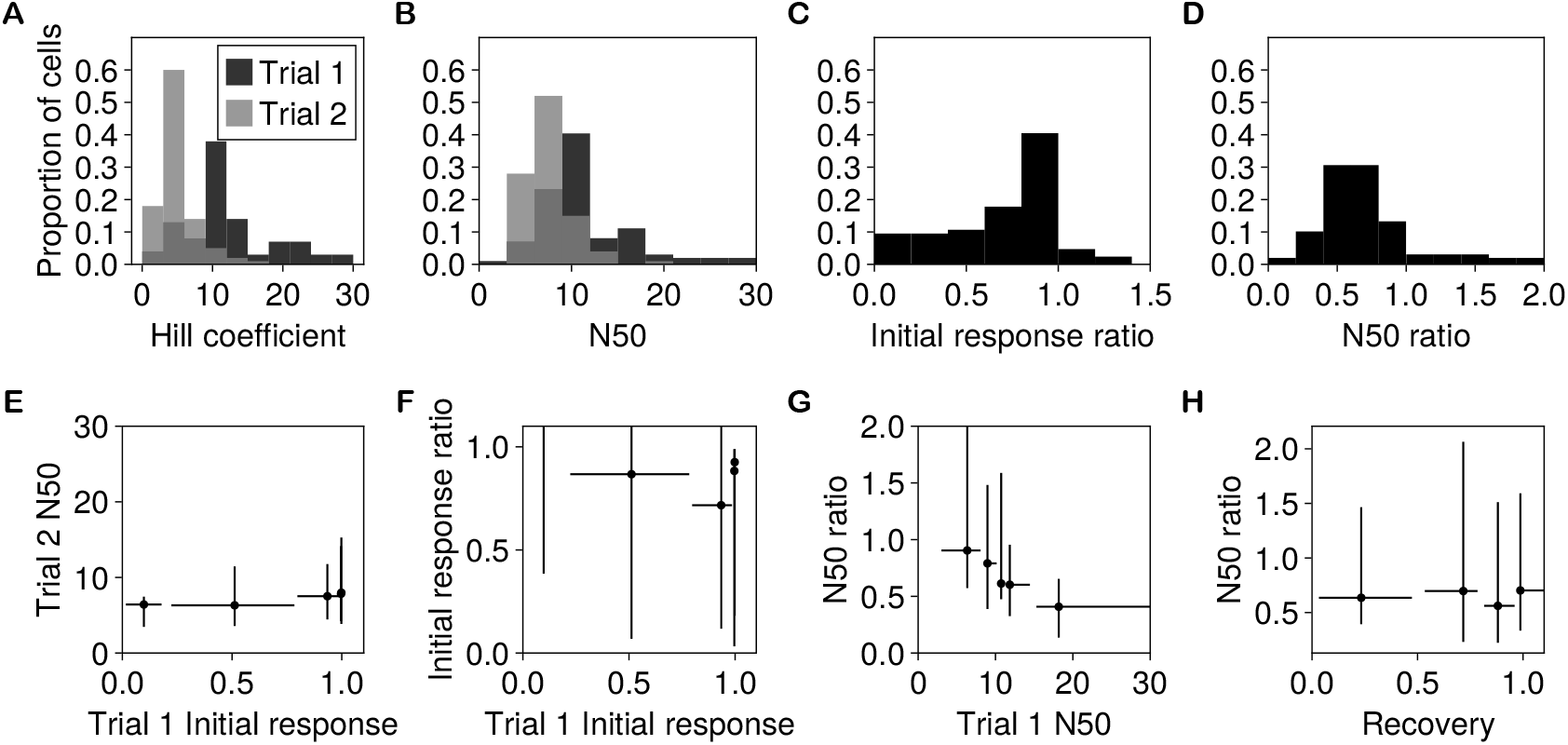
Quantifying relationships between response-profile parameters across cells. **(A)** The posterior for the Hill coefficient is centered far above 1, supporting switch-like habituation at the single-cell level for this 1 min ISI condition. **(B)** Distribution of N50 values across cells for the two trials. **(C)** Distribution of initial response ratios across cells, showing heterogeneous recovery. **(D)** Most cells have a lower N50 in the second trial, indicated by a ratio below 1. (E–H) Scatter plots showing pairwise relationships between inferred parameters across cells. Cells are binned into quintiles by the x-axis parameter; points show within-bin medians; error bars show the central 95% within-bin quantile range of the cell-level posterior median estimates. **(E)** Relationship between trial-1 initial response probability and trial-2 N50. **(F)** Relationship between trial-1 initial response probability and recovery ratio. **(G)** Relationship between trial-1 N50 and N50 ratio. **(H)** Relationship between recovery ratio and N50 ratio.

The N50 parameter indicates the stimulus number at which the response curve has reached 50% of its initial value. Since the population-level N50 parameters control a skewed Gamma distribution, we report our estimate of the median of the resulting distribution across samples. The population-level estimate suggests that cells in this condition take about 11 stimuli to reach half-max (median 11.0, 95% CI 9.4–12.7). The hierarchical model allows us to quantify the heterogeneity across cells that produces this gradual population-level curve. The distribution of the N50 parameter across cells is right-skewed and has a median standard deviation of 6.6 (95% CI 4.7–9.1). In the posterior, 95% of cells cross half-max between 3 stimuli (median 2.9, 95% CI 2.3–4.8) and 27 stimuli (median 27.1, 95% CI 21.7–35.4), pointing to a high degree of variability across cells. Figure 3B shows the corresponding distribution of cell-level N50 estimates.

We filtered the data to control for an equal population mean response for each condition fixed at the mean response across all conditions (0.73). The inferred initial response for this condition had a median of 0.89 (95% CI 0.77–0.97) and a mean of 0.70 (95% CI 0.62–0.77). Cells had a wide distribution of initial responses, with 95% in a window of 0.15–1.0. The median deviates from the mean because this distribution is left-skewed. We can quantify recovery by looking at the ratio between the initial response for trial 2 and trial 1. The median estimate for the ratio of the medians of the two distributions is 0.70 (95% CI 0.52–0.91), showing that for this 1 min ISI, 1 hr ITI condition, the cells have not fully recovered to their baseline response. Figure 3C shows a distribution of median estimates of the recovery ratio for each cell that is consistent with this result. Some cells have a greater initial response in the second trial due to stochasticity; if there were no habituation or recovery, this distribution should be symmetric around 1. The passive recovery period is a window into the decay rate of the process governing habituation, suggesting a time constant for the putative leaky integrator that is consistent with these values.

The distribution of the ratio between trial-2 and trial-1 N50 values is one measure of potentiation. Since the N50 parameter reflects half-max, it is decoupled from the initial response value; the N50 parameter of the second trial reflects the number of stimuli required to reach half of the initial response of the second trial, with partial recovery, rather than half of the initial response of the first trial. The median ratio of the medians of the N50 parameters is 0.64 (95% CI 0.49–0.83), demonstrating that cells reach half-max in 36% fewer stimuli despite starting at a lower initial value. This indicates faster habituation in the second trial with a 1 min ISI and 1 hr ITI. Nevertheless, some cells take longer to reach half-max in the second trial: 95% of cells fall in a window of 0.18–2.07.

We can take advantage of cell-to-cell heterogeneity to compare response parameters and understand the relationships between the underlying processes as a form of “natural experiment.” We computed the correlation between pairs of parameters across individuals for each posterior sample, thereby obtaining a posterior distribution over parameter coupling. Under the assumption of a common dynamical architecture underlying habituation across cells, a correlation suggests shared rate constants influencing the parameters in conjunction.

We find a strong negative relationship between the N50 of the first trial and the N50 ratio between the first and second trials (median correlation - 0.51, 95% CI - 0.64 to - 0.38; Figure 3G). This suggests that the degree of potentiation is coupled to the habituation rate in the first trial: cells that take longer to habituate initially tend to show a larger relative acceleration in the second trial. This is consistent with a hierarchical cascade model in which a slow second-order memory trace is charged by the first-order habituation process during the first trial.

We can interpret the initial response to the first stimulus in trial 1 as a reflection of the cell’s overall responsiveness, assuming that this difference is endogenous before stimulation begins. Each cell has a variable propensity to respond to stimuli of a particular strength. ***Wood*** (***1969***) conducted an elegant study to determine whether habituation depends on the number of contractions or the number of stimuli. We can indirectly test this result in our framework by asking how baseline responsiveness relates to the heterogeneity of habituation time. If the negative feedback mechanism were downstream of the action potential or contraction apparatus, we might expect a more responsive cell to have a lower N50 value: it starts at a higher initial response probability and is therefore more likely to contract to stimuli, so each contraction could drive a greater decrease in response probability. We observe instead that the initial response is not significantly correlated with N50 (median 0.07, 95% CI - 0.06 to 0.22), suggesting that habituation time is largely disconnected from baseline responsiveness. This is consistent with Wood’s finding that it was the number and pattern of stimulation rather than the number and pattern of contractions that controls the degree of habituation. On the other hand, there is a negative correlation between initial response and recovery ratio (median - 0.32, 95% CI - 0.53 to - 0.19; Figure 3F), suggesting that more responsive cells recover more slowly. Lastly, we find no significant correlation between recovery and potentiation (median - 0.07, 95% CI - 0.15 to 0.08; Figure 3H). This is consistent with the decay constants for the two orders of memory being at least partially independent.

### Habituation is frequency-sensitive in absolute time

Frequency sensitivity describes the property wherein habituation and recovery are faster for higher-frequency stimuli. In ***Wood*** (***1969***), *Stentor* showed partial frequency sensitivity: habituation to stimuli delivered at 1 min ISI is faster than habituation to 2 and 3 min ISI stimuli, with the latter two conditions being roughly equal. In our apparatus, 1 min is a practical lower bound on the ISI due to the re-extension time after contraction, which would lead to double counting for shorter intervals. Nevertheless, ***Rajan et al. (2023***) found that a 1.2 s ISI stimulus train leads to faster habituation than a 1 min ISI stimulus train, which is consistent with frequency sensitivity.

Neither study examined recovery as a function of stimulus frequency, which is important mech-anistically. Computational models of habituation show that although a single leaky integrator mechanism is sufficient to explain faster learning to higher-frequency stimuli by virtue of accumulating a greater signal within a given time, it predicts slower recovery after stronger accumulation when the decay time constant is fixed. In contrast, a cascade of leaky integrators with separate time constants can show both faster learning and faster recovery for higher-frequency stimuli (***Staddon and Higa, 1996***). Intuitively, each unit in the cascade is tuned to a particular frequency tied to its time constant.

Figure 4 shows the aggregated response of the cells in our study for the first trial of each ISI. *Stentor* habituation does not appear strongly frequency-sensitive upon simple examination of the population response. Across all three ISIs, we observe similar learning rates and asymptotic population means when response is plotted by stimulus number. One possibility is that there is no decay of the leaky integrator or that the time constant is very long, leading to perfect integration (***Drew and Abbott, 2006***). This would lead to identical habituation curves across all frequencies as a function of stimulus number; essentially, time would not be a factor. However, in the absence of some other competing process, this would also mean that cells take an indefinitely long period to recover or do not recover to baseline at all. Another possibility is that *Stentor* habituation is time-sensitive, but that *Stentor* have mechanisms that accommodate this three-fold difference in frequencies almost perfectly.

**Figure 4.**
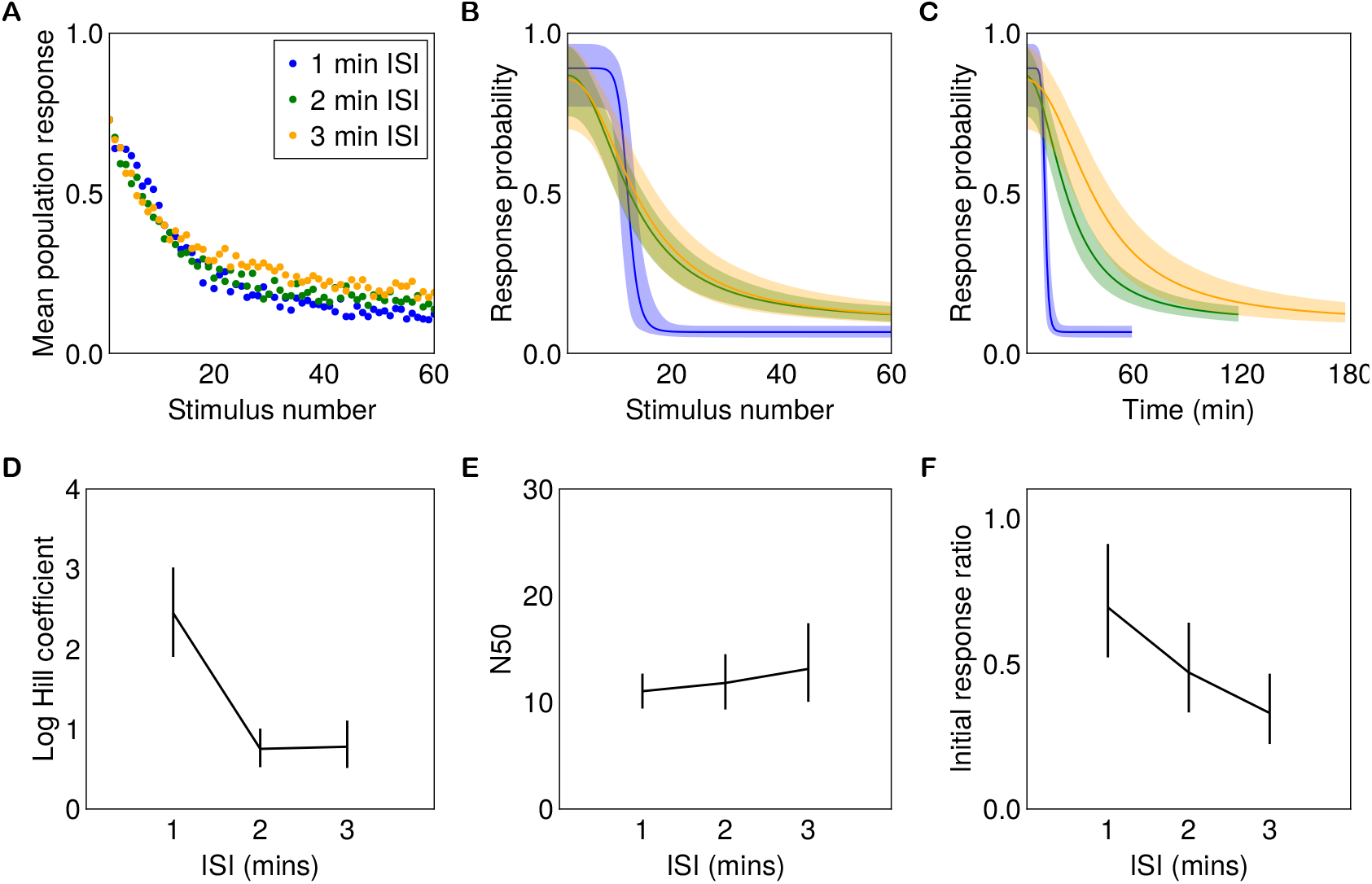
Frequency sensitivity of *Stentor* habituation. **(A)** We varied the ISI from 1 min to 3 min. As a function of stimulus number, the population mean response looks similar across ISIs. **(B)** Inferred trial-1 response curves plotted as a function of stimulus number. **(C)** The same inferred curves plotted as a function of time; shorter ISI leads to faster habituation in absolute time. **(D)** The Hill coefficient is higher for the 1 min ISI condition than for the 2 or 3 min conditions, suggesting a switchier habituation profile at the single-cell level. Error bars show 95% credible intervals. **(E)** Median half-max values are similar across ISIs when measured in stimulus-number units. **(F)** Recovery is frequency-sensitive: recovery ratios after a 1 hr ITI decrease with ISI.

Nevertheless, the single-cell results point to differences across frequencies. The high-frequency condition is more switch-like: the first trials of the 1 min ISI conditions generally have higher Hill coefficients (medians 11.6, 5.0, 3.0, 8.1) compared to the 2 min (medians 2.1, 2.2, 3.7, 2.3) and 3 min (medians 2.2, 3.4, 1.9, 1.4) ISI conditions, suggesting higher cooperativity (Figure 4D). For a fixed N50, this makes the initial decrease of response probability slower as a function of stimulus number, but the curve eventually catches up and exceeds the habituation rate of the longer ISI conditions. We successfully control for the initial response across ISIs for the 1 hr ITI conditions shown in Figure 4B–C, yielding median values of 0.89, 0.87, 0.86 (95% CIs 0.77–0.97, 0.74–0.96, and 0.70–0.96). We note, however, that there is variability across trials with the same ISI, reflecting the challenge of controlling response curves across experiments.

A key distinction here is defining habituation rate as a function of stimulus number versus time. It is impossible to hold both the number of stimuli delivered and the total duration of the trial constant when changing the ISI. Viewed as a function of time, the single-cell curves are ordered by ISI, with the shortest ISI decreasing the fastest (Figure 4C). Viewed as a function of stimulus number, the trial-1 curves for all conditions have similar N50 medians of 11.0 (95% CI 9.4–12.7), 11.8 (95% CI 9.3–14.5), and 13.1 (95% CI 10.1–17.4) for the 1, 2, and 3 min conditions, respectively (Figure 4E). The N50 parameters are not credibly different across conditions. However, habituation is frequency-sensitive as a function of time, which is expected given the greater total stimulus delivered in a given unit of time. The fact that all three curves are relatively close despite the 1 min ISI condition receiving substantially more input suggests that absolute-time effects may also be at play, perhaps due to the characteristic timescale of the underlying process. For example, it might take a minimum amount of time for a kinase to become active. There is also no significant difference between the asymptotes of the curves.

To further narrow down the potential models, we investigated the recovery profiles for each ISI by querying the response probabilities of the cells after the 1 hr recovery period. Consistent with frequency sensitivity, we find that *Stentor* take longer to recover after longer-ISI protocols. With a recovery duration of 1 hr, as shown in Figure 4F, the 1, 2, and 3 min ISI conditions have median recovery ratios of 0.70 (95% CI 0.52–0.91), 0.47 (95% CI 0.33–0.64), and 0.33 (95% CI 0.23–0.47), respectively. This suggests that the mechanism underlying habituation to longer-ISI stimuli operates at a corresponding timescale. This could be due to varied temporal tuning of the relevant molecular species or to distinct molecular species.

### Varying the ITI demonstrates the effect and decay of potentiation

In addition to the ISI, we varied the recovery period (ITI) between the first and second trials to test how this affected learning in the latter. We gathered data for every combination of ISIs (1–3 min) and ITIs (1, 2, 3, and 5 hr). This set of parameters was chosen so that there was one condition where the ITI was as long as the duration of the first trial for each ISI, i.e., the ITI was 60 times longer than the ISI. This yielded a total of twelve conditions, the results of which are shown in Figure 5. Longer experiments are challenging to complete, which prevented us from doing comparable studies at 5 min ISI.

**Figure 5.**
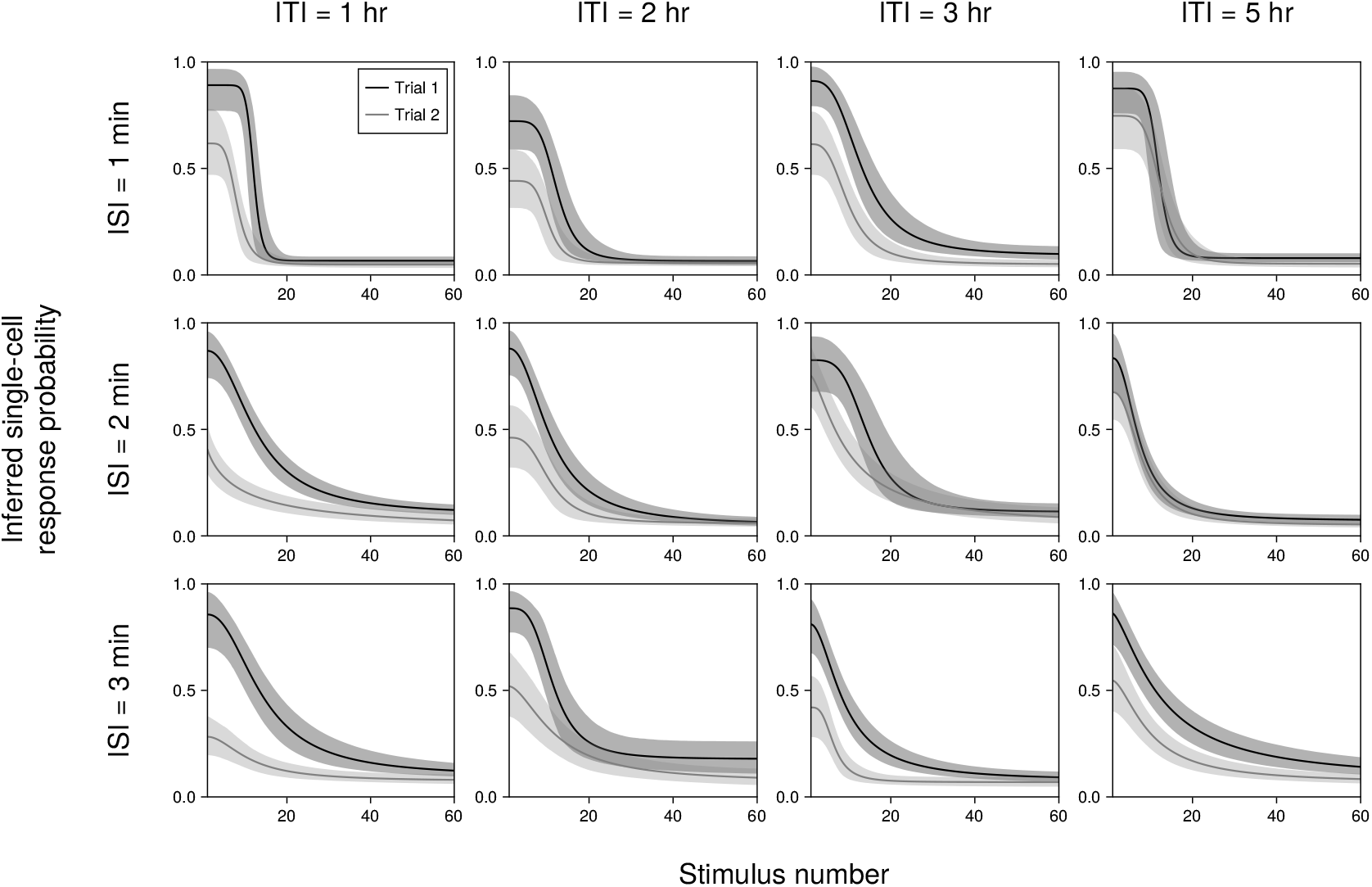
Inferred single-cell curves plotted with population parameters. Each axis displays the inferred single-cell curves for a condition with a particular ISI (1, 2, 3 minutes) and ITI (1, 2, 3, 5 hours). Bands show curves corresponding to the 95% credible interval.

When the ITI is too short, the response has not recovered to baseline. Thus, the “first-order” memory by our definition, i.e., the diminution of response probability, still persists. As a result, it is challenging to observe the second-order memory directly. Although it is clear in all the 1 hr ITI experiments that the response in the second trial is lower than the response to the corresponding stimulus in trial 1, it is unclear from response level alone whether the apparent difference in habituation is due to different initial response values for each trial or a true difference in learning rate.

### Decoupling response level and learning rate

To analyze potentiation with stronger foundations, we return to its explicit definition: the learning rate can be higher in the second trial despite the cell having the same response probability as in the first trial. The direct way to examine this is to plot the derivative of the response probability against the response probability itself, which is effectively the inferred phase portrait of the underlying dynamical system:

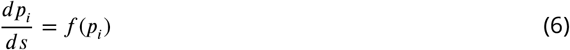

The function *f* can be estimated empirically by taking the derivative of the response curve for any given cell and plotting it against the response probability. ***Figure 6***—***figure Supplement 1*** illustrates how a learning curve can be translated into a phase portrait. This obviates the issue wherein partial recovery can falsely suggest potentiation even though the learning dynamics are unchanged. ***Figure 6***—***figure Supplement 2*** illustrates this using a toy example.

**Figure 6.**
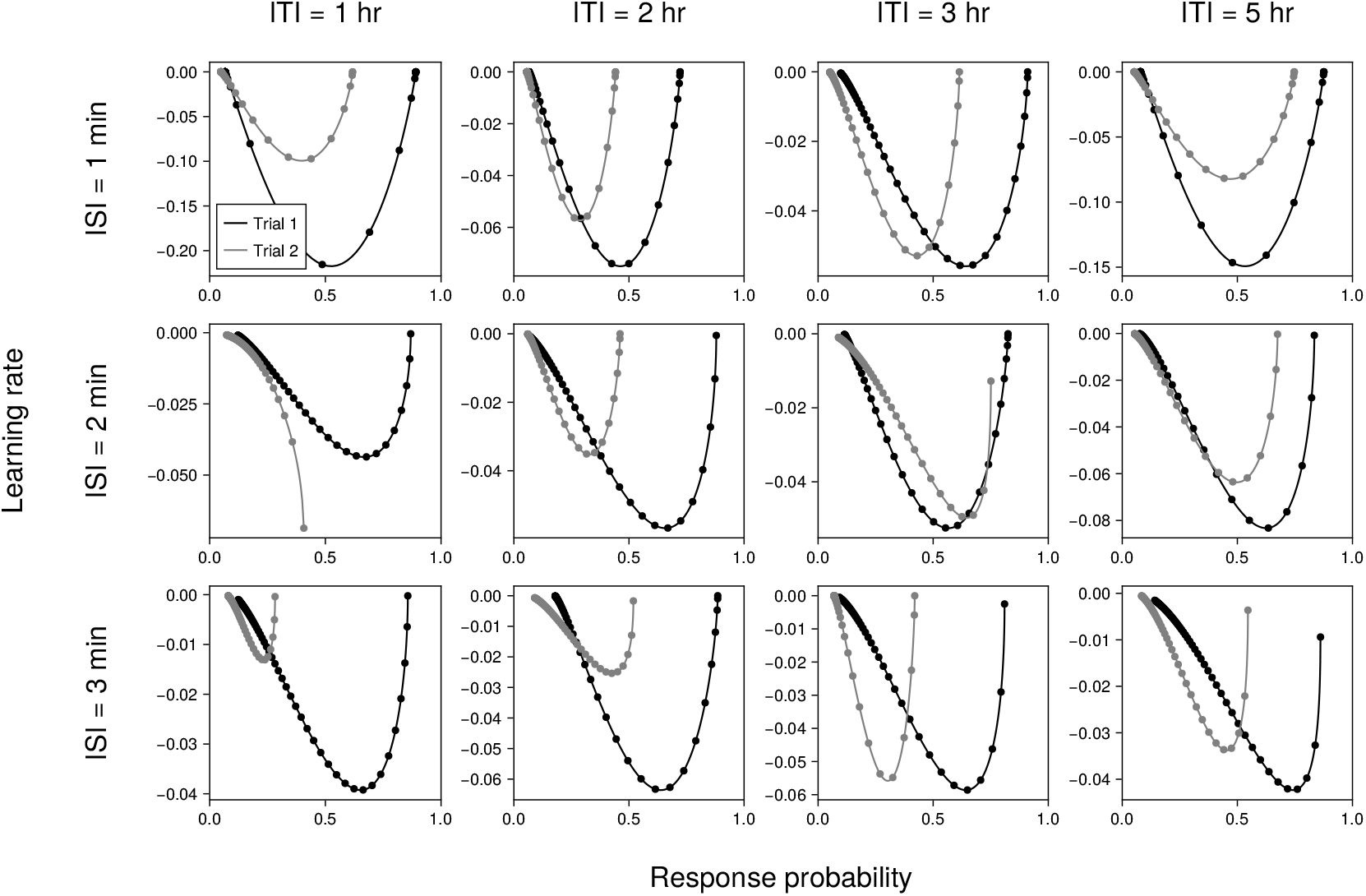
Inferred phase portraits. Each axis displays the phase portraits computed using the inferred single-cell curves for a condition with a particular ISI (1, 2, 3 minutes) and ITI (1, 2, 3, 5 hours). Scatter points show progression by stimulus number.

Figure 6 shows the phase-portrait equivalents for the curves in Figure 5. The phase portraits deviate between trial 1 and trial 2 across all 12 conditions, suggesting the presence of additional state variables that control the dynamics of habituation.

### Formally quantifying potentiation

Potentiation is defined as faster learning in the second trial compared to the first (***Rankin et al., 2009***). However, it is not always clear what “faster” means. Figure 7 presents three ways of mathematically formalizing this definition. Figure 7A shows N50, the half-max time calculated directly from the inferred Hill curve parameters, as in Figure 3D. In Figure 7B, we compute the derivative of the habituation profile, which we refer to as the learning rate, and normalize it by the initial response level. We account for noise by averaging across the first five stimuli. Finally, we compute the cumulative absolute difference between the phase portraits for trial 1 and trial 2 as a measure of deviation between the dynamics of the two trials (Figure 7C).

**Figure 7.**
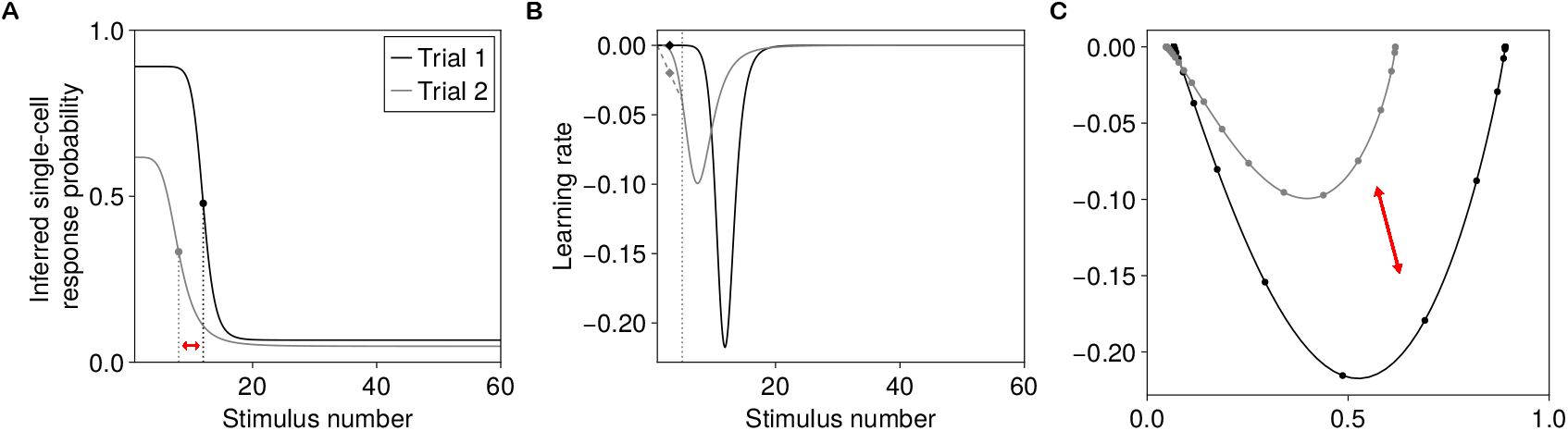
Quantifying potentiation across two trials. **(A)** Ratio between N50 (half-max) values. **(B)** Ratio between initial learning rates, discretized by averaging across the first five stimuli. **(C)** Cumulative distance between phase portraits captures deviation in dynamics.

We then investigate how recovery (initial response ratio) and potentiation vary across combinations of stimulus parameters. Figure 8A shows the recovery profiles for the 1, 2, and 3 min ISI conditions across the range of rest durations (ITIs). We see an ordering in the magnitude of recovery at the 1 hr recovery period, as shown in Figure 4F. The initial response ratios for the 1 min and 3 min ISI conditions are significantly different, with a median ratio of 0.48 (95% CI 0.30–0.75). However, with a 5 hr recovery period, the median ratio is higher at 0.75 (95% CI 0.50–1.11). Thus, the longer-ISI condition is slower to recover than the shorter-ISI condition. This establishes that recovery is frequency-sensitive. The recovery profile is a function of absolute time, not number of stimuli. In sum, our data suggest that *Stentor* habituation and recovery are frequency-sensitive in terms of absolute time but not in terms of stimulus number.

**Figure 8.**
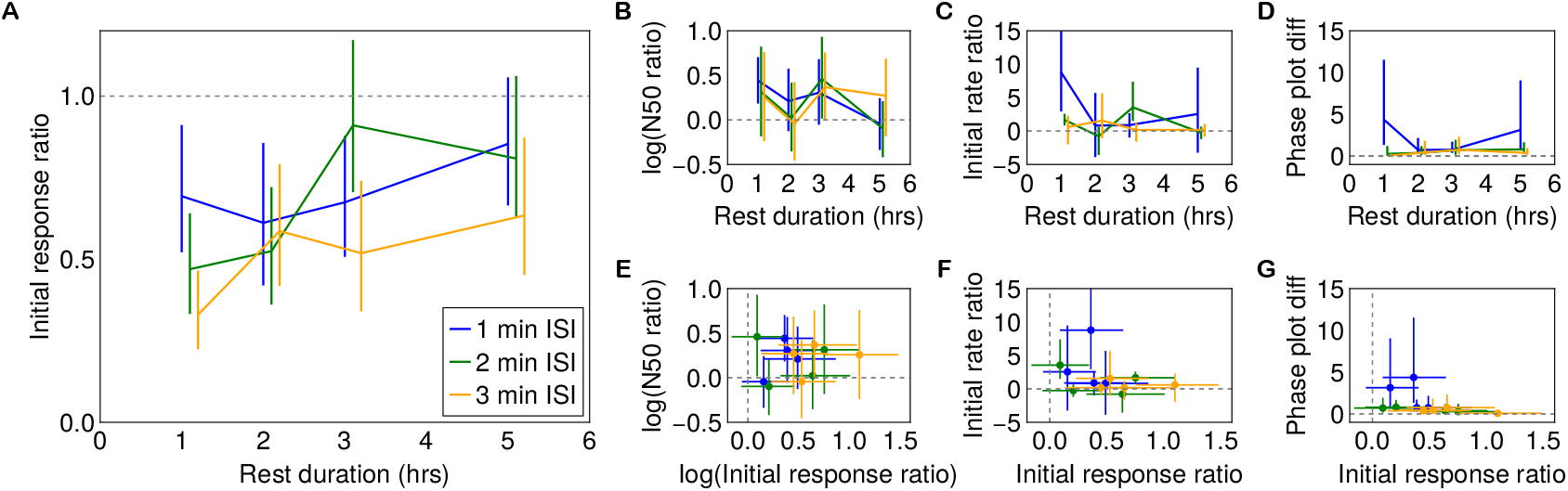
Quantifying potentiation across ITIs. **(A)** Initial response recovery curves for each ISI condition across all recovery periods with 95% credible intervals. **(B)** Log N50 advantage decreases toward zero with increasing ITI. **(C)** Initial learning-rate ratio similarly decays with rest duration. **(D)** Cumulative phase-portrait difference decays with rest duration. **(E–G)** Recovery (x-axis) plotted against each potentiation metric (y-axis) for all ISI/ITI combinations. The spread across ISI conditions at similar recovery levels suggests partial decoupling of the recovery and potentiation processes.

We next ask whether potentiation, like recovery, decays with the rest duration. Figures 8B–D plot the three potentiation metrics—the log N50 advantage, the initial learning-rate ratio, and the cumulative phase-portrait difference—as functions of ITI for each ISI condition. As the ITI increases, the log N50 advantage converges toward zero (Figure 8B), meaning that the advantage in half-max time for the second trial diminishes. Similarly, the initial learning-rate ratio (Figure 8C) and phase-portrait difference (Figure 8D) decrease toward zero, indicating that the dynamics of trial 2 converge to those of trial 1 as the rest duration lengthens. This is consistent with the gradual decay of a second-order memory trace that mediates potentiation.

Finally, Figures 8E–G plot recovery (log initial response ratio) against each potentiation metric, with each point representing an ISI/ITI condition. If recovery and potentiation were controlled by the same process, we would expect a tight, monotonic relationship between the two. Instead, we observe that the relationship between recovery and potentiation varies across conditions: for a given level of recovery, different ISI conditions can have different potentiation magnitudes. This suggests that the first-order memory governing habituation (and its recovery) and the second-order memory governing potentiation decay on distinct timescales and are at least partially decoupled, consistent with a serial cascade architecture in which a slow integrator encodes potentiation downstream of the faster habituation process.

## Discussion

We used habituation as a window into the learning capabilities of a single-celled organism. Our study builds on classic experiments by ***Wood*** (***1969***) and recent experiments by ***Rajan and Marshall (2025***) showing that *Stentor* exhibits both habituation and potentiation. We systematically varied stimulus frequency and recovery duration across 12 conditions, collected single-cell contraction data at scale, and developed a hierarchical Bayesian framework to infer latent habituation curves for individual cells. By inferring latent response probabilities at the single-cell level, we asked how these phenomena vary across cells and across stimulus protocols, rather than treating the population mean as the primary object of analysis. A phase-portrait analysis allowed us to decouple response level from learning rate, providing a stringent test for potentiation. Our key findings are that (i) habituation is approximately invariant to frequency when measured in stimulus-number units but frequency-sensitive in absolute time, (ii) recovery is also frequency-sensitive, with longer-ISI protocols producing slower recovery, (iii) single-cell habituation is switch-like at higher frequencies but more gradual at lower frequencies, (iv) potentiation is present across conditions and decays with the intertrial interval, and (v) recovery and potentiation are largely decoupled.

These results suggest that memory in *Stentor* is not captured by a single recovery clock. We argue that, in habituation, long-term memory should be understood not only in terms of absolute duration, but also in terms of how underlying memory processes interact across timescales. If habituation is a useful model for studying plasticity, then potentiation can serve as a model for studying metaplasticity: the plasticity of plasticity (***Abraham and Bear, 1996***). Our results support the existence of distinct but coupled memory processes operating on different timescales: a faster process that changes response probability within a stimulus sequence and a slower process that changes how rapidly the first process is engaged in a later sequence.

This separation of timescales is clearest in the contrast between habituation and recovery. The degree of habituation is approximately invariant to frequency when viewed as a function of stimulus number, suggesting that the number of stimulus presentations is a strong determinant of response decrement. Recovery, by contrast, can only be measured in units of time and varies with ISI. This dissociation implies that the underlying processes have different time constants, akin to the leaky integrator cascade models presented by ***Staddon and Higa*** (***1996***). Our analysis suggests that the processes governing recovery and potentiation are largely decoupled, with low or no correlations between the parameters governing them across cells. Thus, recovery of response probability does not necessarily imply erasure of the latent state that controls future learning rate. However, our data are too noisy to determine the timescale of this putative latent mechanism.

The single-cell fits further suggest that the mechanism producing response decrement may depend on both stimulus count and absolute time. Our inference of the Hill coefficient suggests that cooperativity is higher for the higher-frequency condition (1 min ISI) than for the lower-frequency conditions. Viewed as a function of stimulus number, the single-cell habituation curve for the higher-frequency condition appears to be slower for the initial stimuli before surpassing the habituation level of the lower-frequency conditions. One possible interpretation is that the process producing the decrement in response has a characteristic activation timescale, such that its effects become more apparent only after several stimuli. Alternatively, this response profile could result from competing habituation and facilitation processes. Intracellular calcium may accumulate during the initial trials, especially with shorter intervals, if the rate of clearing is slow enough. Presynaptic facilitation as a result of calcium accumulation has been shown to be responsible for sensitization in Aplysia (***Pinsker et al., 1973***; ***Castellucci and Kandel, 1976***).

Our results on potentiation are consistent with work in mammalian neuron-like cells (***Cheever and Koshland, 1992***), one of the few studies to explicitly examine the rate constants involved in putative receptor modification in a negative feedback circuit. However, the duration of that protocol is limited to approximately 1 hr with ∼10 stimulus applications separated by 4 min. *Stentor*, on the other hand, show habituation and modulation of learning rate over much longer durations.

Massed versus spaced training paradigms in neuroscience (***Smolen et al., 2016***), which have also been demonstrated in non-neural cells (***Kukushkin et al., 2024***), typically study changes in learning rate by varying the spacing of a small number of stimuli (∼ 10). *Stentor* habituation demonstrates that unicellular organisms can show varied responses to more complex learning protocols over longer durations of time, due to coupling between processes operating across a wide range of timescales: contraction (action potential) takes milliseconds, relaxation takes tens of seconds, habituation and recovery take tens of minutes or longer, and potentiation lasts for hours, at least until full recovery or beyond. The fastest process, mediated by mechanoreceptor currents, and the slowest process, mediated by a putative modification of the mechanoreceptor (***Wood, 1988***), are separated by over five orders of magnitude in time.

Extrapolating from the frequency-invariance result suggests that *Stentor* can habituate to stimulus series with much longer ISIs. Indeed, in unpublished data, we find that habituation occurs at 10 min and even 30 min ISIs. However, experimental limitations constrain the number of stimuli we can present at longer ISIs, and thus we can neither observe the asymptotic degree of habituation nor measure the recovery profile. Future experiments with longer timescales will be needed to determine whether the same stimulus-count dependence persists under sparser stimulation.

Another limitation of our study is that our probabilistic model does not have shared parameters across conditions due to computational constraints. Shared parameters would have allowed us to control for overall responsiveness, which should be identical across conditions, and for the first-trial response profile, which should be identical across conditions with a given ISI. Our data are therefore too noisy to make strong claims about higher-order relationships between stimulus parameters and response profiles.

Our experimental and mathematical analysis of potentiation also suggests a direct test of the proposed second-order memory trace. If we pause a habituation trial and perturb the response level back to baseline, the cells should still show faster learning when the trial is continued, provided that the second-order memory trace remains active. Such perturbation experiments would test whether recovery of response probability and erasure of the potentiating memory can be dissociated within the same cell.

Although several computational models of habituation can reproduce various hallmark characteristics, few have confronted the vast range of timescales over which the mechanisms underlying habituation operate. Even if a simple two-unit model can show potentiation, it is insufficient to demonstrate how these mechanisms are coupled. This is important mechanistically because a larger separation of timescales might necessitate more layers of this hierarchy. Future modeling work needs to contend with the range of timescales at which the various processes operate and, consequently, with constraints on the coupling between them.

Studying habituation in the ancient single-celled organism *Stentor* allows us to understand learning and memory across the tree of life. Our focus on potentiation points to common computational motifs and intracellular mechanisms not only for plasticity, but also for metaplasticity. This serves as a useful tool for understanding molecular computation in single cells more broadly, including neurons (***Bhalla, 2014***).

## Materials and Methods

### Cell culture

*Stentor coeruleus* were obtained from Carolina Biological (catalog no. 131598) and maintained in filtered spring water (Carolina Biological, catalog no. 132450) for a minimum of two weeks before experimentation. Cultures were kept in Pyrex bowls in dark drawers at 22° C when not in use.

*Chlamydomonas reinhardtii* (Chlamydomonas Resource Center, CC-125) were grown on 1.5% agar plates prepared with TAP medium (Phytotechlabs, T8224) under continuous bright light. *Stentor* cultures were fed with *Chlamydomonas* every 4–5 days. For each feeding, five loopfuls of *Chlamydomonas* were transferred using a sterile inoculating loop into 50 mL of filtered spring water. Large aggregates were broken up by gentle pipetting and shaking before feeding.

Both *Stentor* and *Chlamydomonas* cultures were inspected weekly for signs of contamination, cell death, or reduced growth. *Stentor* cultures were transferred to fresh spring water monthly to maintain culture health and minimize bacterial overgrowth.

### Experimental apparatus

Each habituation device housed a solenoid (Adafruit Mini Push-Pull 5V) positioned beneath the center of a 35 mm Petri dish coated with poly-D-lysine (Mattek) containing *Stentor*. Solenoids were connected to a programmable power supply (KORAD KA6003P) and controlled via a Raspberry Pi Pico running MicroPython. Upon activation, the solenoid piston was displaced upward by 2.5 mm, contacted the dish to deliver a mechanical tap, and then returned to its resting position. Stimulus strength was controlled by adjusting the supply voltage, and stimulus timing was programmatically specified to allow precise control over the ISI and ITI. Each experimental device was equipped with an overhead-mounted camera (FLIR 11-515) and a 24-bit Neopixel LED ring for illumination. The LED ring was covered with diffusion paper to minimize glare and provide uniform lighting across the field of view. Videos were recorded at 10 frames per second.

For each experiment, 400 *µ*L of *Stentor* culture was transferred to 2300 *µ*L of filtered water in the Petri dish. Each run was preceded by an incubation time of 8 hours to allow cells to recover from the mechanical stress of handling. The incubation concluded with a 2-hour window under illumination to acclimate the cells to imaging conditions.

Experiments were conducted inside a custom-built, light-shielded acrylic enclosure mounted on a vibration isolation platform (ThorLabs, PTT600600). Additional vibration damping was provided by four sorbothane feet attached to an optical breadboard (ThorLabs, AV6).

### Video annotation

Behavioral recordings were synchronized with stimulus delivery. For each stimulus, video was captured starting 1 s before solenoid activation and continuing for 1 s following the stimulus. This allowed quantification of cell morphology and contraction behavior immediately before and after mechanical stimulation.

Custom video annotation tools were developed to score contraction responses at the single-cell level. Cells were manually labeled as contracting or non-contracting in response to each stimulus across the full experimental protocol. This annotation enabled tracking of individual cell responses across both stimulus sequences and quantitative comparison of response dynamics. Only cells that remained anchored through the entire run (trial 1, ITI, trial 2) were used for analysis, consistent with Wood’s methods. Additionally, we excluded cells that underwent mitosis or conjugation, or made contact with neighboring cells.

### Probabilistic modeling

To quantitatively characterize habituation and potentiation at the single-cell level, we developed a probabilistic model of contraction responses using Bayesian inference. Modeling was implemented in Julia using the probabilistic programming package Turing.jl.

Each cell’s response to a stimulus was treated as a binary random variable indicating contraction or non-contraction. We subsampled the contraction responses to make the initial population mean response for the first trial of each condition match the mean across all conditions. The probability of contraction was modeled as a function of stimulus number within a sequence, allowing the response probability to decrease over repeated stimulation. The same single-trial model was fit separately to the first and second stimulus sequences, enabling direct comparison of habituation dynamics across sequences. The probabilistic model is shown in Box 1. At the population level, cell-specific parameters were drawn from shared prior distributions, yielding a hierarchical model that captured both inter-cell variability and common structure across the population.

Posterior inference was performed using No-U-Turn Sampling (NUTS), with automated mass matrix estimation, as implemented in Turing. We collected 2500 samples after a burn-in period of 2500. Convergence was assessed using standard diagnostics, including 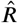 (criterion <1.03) and effective sample size (ESS). Most parameters had an 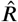 in the range 0.99–1.01 and an ESS in the range 500–1000. The resulting Markov Chain Monte Carlo (MCMC) chain for the posterior distribution was used for subsequent analysis as described in the main text. Derived quantities at the Probabilistic model

**Hyperpriors for trial***r*

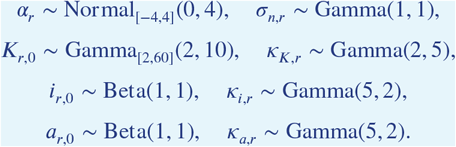

Subscripted intervals indicate truncated distributions.

**Cell-level parameters for cell** *j*

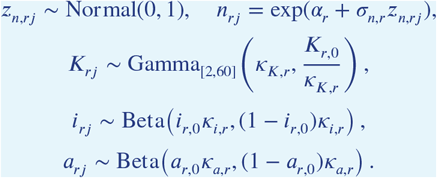

Observations for stimulus *s*

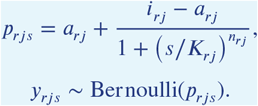

population level (differences, ratios) and at the inter-individual level (correlations) were computed per draw to obtain the corresponding distributions.

## Acknowledgment

We thank Deepa Rajan and Wallace Marshall for help with our *Stentor* culture, Matt Laudon for help with our *Chlamydomonas* culture, and John Vastola, Yue Miao, Zachary Kelso, Madeleine Snyder, and Jeremy Gunawardena for helpful discussions. This work was supported by a Polymath Award from Schmidt Sciences and a grant from the Air Force Office of Scientific Research (FA9550-22-1-0345).

---

**Figure Figure 6—figure supplement 1.**
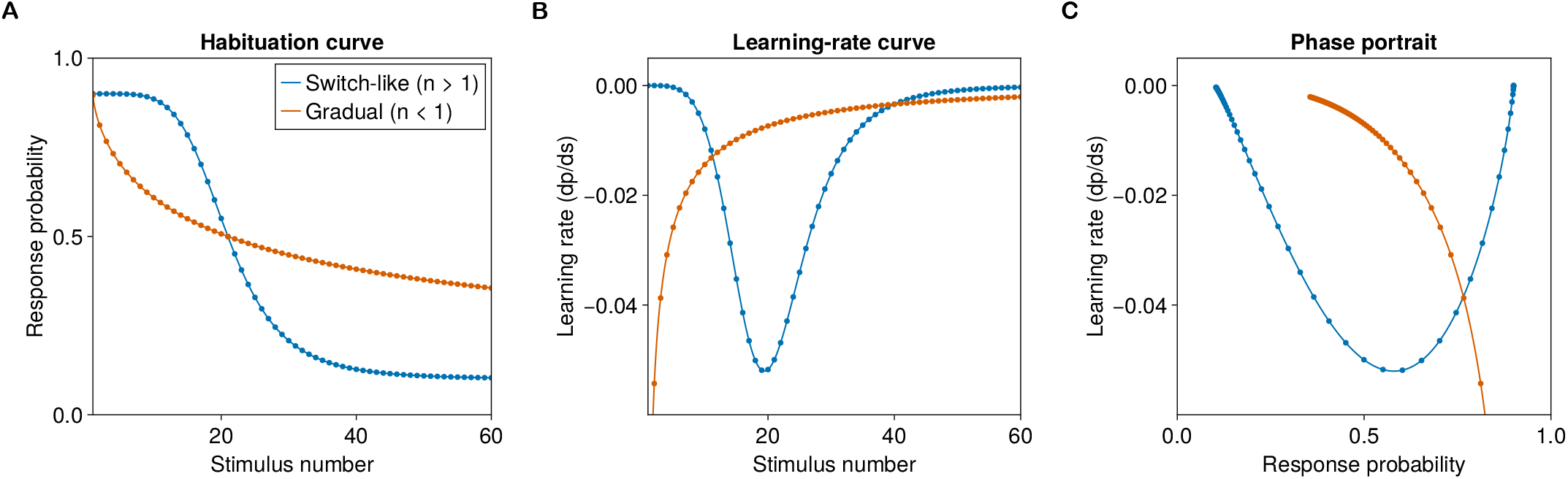
Translating habituation curves to phase portraits. **(A)**Example habituation curves for a switch-like Hill function (*n >* 1) and a gradual Hill function (*n <* 1).**(B)** Taking the derivative of each curve with respect to stimulus number gives the learning-rate curve. **(C)** Plotting the learning rate against response probability yields the phase portrait. The gradual curve is monotonic and begins with a nonzero learning rate, whereas the switch-like curve is U-shaped and begins close to zero. Dots indicate stimulus number and become denser as the curve asymptotes.

**Figure 6—figure supplement 2.**
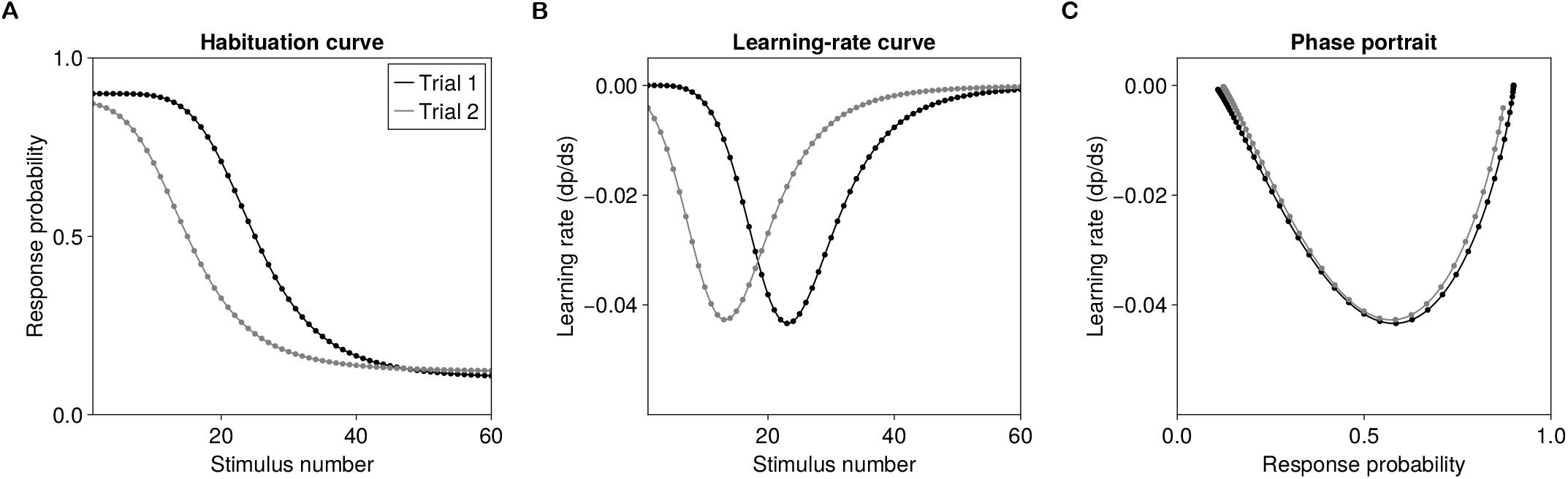
Partial recovery can preserve the phase portrait. **(A)** Example trial-1 and trial-2 habituation curves. The trial-2 curve starts from a partially recovered response level, with a small perturbation to the curve parameters (for illustrative purposes). The halfmax stimulus is 24.88 for trial 1 and 15.05 for trial 2. **(B)** The corresponding learning-rate curves differ when plotted against stimulus number. **(C)** Plotting learning rate against response probability yields similar phase portraits, showing that visually different habituation curves can arise from nearly identical response-level dynamics. Dots indicate stimulus number and become denser as the curves asymptote.

